# The dimerisable Cre recombinase allows conditional genome editing in the mosquito stages of *Plasmodium berghei*

**DOI:** 10.1101/2020.07.13.200402

**Authors:** Priyanka Fernandes, Sylvie Briquet, Delphine Patarot, Manon Loubens, Bénédicte Hoareau-Coudert, Olivier Silvie

## Abstract

Asexual blood stages of the malaria parasite are readily amenable to genetic modification via homologous recombination, allowing functional studies of parasite genes that are not essential in this part of the life cycle. However, conventional reverse genetics cannot be applied for the functional analysis of genes that are essential during asexual blood-stage replication. Various strategies have been developed for conditional mutagenesis of *Plasmodium*, including recombinase-based gene deletion, regulatable promoters, and mRNA or protein destabilization systems. Among these, the dimerisable Cre (DiCre) recombinase system has emerged as a powerful approach for conditional gene targeting in *P. falciparum*. In this system, the bacteriophage Cre is expressed in the form of two separate, enzymatically inactive polypeptides, each fused to a different rapamycin-binding protein. Rapamycin-induced heterodimerization of the two components restores recombinase activity. We have implemented the DiCre system in the rodent malaria parasite *P. berghei*, and show that rapamycin-induced excision of floxed DNA sequences can be achieved with very high efficiency in both mammalian and mosquito parasite stages. This tool can be used to investigate the function of essential genes not only in asexual blood stages, but also in other parts of the malaria parasite life cycle.

## INTRODUCTION

The life cycle of *Plasmodium* parasites is complex, and involves several stages of development and differentiation within the vertebrate host and *Anopheles* vector. In mammalians, infection begins when motile forms of the parasite known as sporozoites are transmitted by infected *Anopheles* mosquitoes to the host. These sporozoites first go through an obligatory intra-hepatic step, where they multiply into thousands of merozoites that are released into the blood stream, where they begin the next round of replication within erythrocytes. Some merozoites develop into gametocytes which fuse to form a motile ookinete in the gut of the mosquito. These ookinetes encyst on the basal side of the midgut wall, where they further differentiate into sporozoites that invade the salivary glands, and are transmitted by the mosquito during the next blood-meal [1].

Experimental genetics have been instrumental in deciphering key aspects of the parasite biology. However, a thorough understanding of the genes that are critical for several stages of development, is lacking. The *Plasmodium* genome is haploid during the intra-erythrocytic stage of development and is thus easily amenable to genetic modification via homologous recombination. Hence most gene modification systems are adapted to targeting genes at the blood-stage of the parasite life cycle. However, as conventional reverse genetic strategies cannot be used to target genes that are essential for blood-stage replication, the functional analysis of these genes in other stages of development remains largely unexplored.

Various strategies have previously been developed for conditional mutagenesis in *Plasmodium*, including recombinase-based gene deletion [2], regulatable promoters [3], and mRNA or protein destabilization systems [4–7]. While most of these strategies are stage-specific, others require a promoter that is expressed by distinct stages of the parasite, thus limiting the investigation of essential genes in multiple developmental stages. In recent years, the dimerisable Cre (DiCre) recombinase system has emerged as an efficient system for conditional gene knockout in *P. falciparum* and other apicomplexans [2,8]. It consists of the expression of the bacteriophage Cre in the form of two separate, enzymatically inactive polypeptides, each fused to a different rapamycin-binding protein (either FKBP12 or FRB) that heterodimerise after the addition of rapamycin [9]. Heterodimerisation of the two components in turn restores recombinase activity, leading to site-specific excision of Lox-flanked DNA sequences and rapid and efficient rapamycin-induced gene deletion [2,8]. We implemented a similar approach in *P. berghei*, a rodent parasite model that is used to study malaria in the laboratory, as its life cycle can be completed by cycling between infected mice and *Anopheles* mosquitoes. We first generated a *P. berghei* parasite line that stably expresses both components of the DiCre cassette, and validated that the resulting parasite line showed no phenotypical defect as compared to the parental strain. We show that a single rapamycin dose administered to mice is sufficient to induce efficient excision of a floxed target DNA sequence *in vivo* in mice, including in transmission stages. By adapting the rapamycin treatment to mosquitoes, we also show that Cre-mediated excision can be achieved in mosquito stages, thus demonstrating the versatility and robustness of the DiCre system to inactivate genes across the parasite life cycle.

## MATERIALS AND METHODS

### Ethics statement

All animal work was conducted in strict accordance with the Directive 2010/63/EU of the European Parliament and Council on the protection of animals used for scientific purposes. Protocols were approved by the Ethical Committee Charles Darwin N°005 (approval #7475-2016110315516522).

### Generation of plasmids for transfection

#### DiCre plasmid

The DiCre plasmid was designed to replace the GFP cassette previously introduced in the *p230p* locus of *P. berghei* ANKA parasites [10] by mCherry, DiCre and TgDHFR/TS cassettes. The DiCre plasmid was generated by assembling five DNA fragments: a mCherry cassette flanked by a 800-kb 3’ fragment of *P. berghei* HSP70 promoter and PbDHFR 3’UTR (DC1); a Cre59 coding sequence followed by PfCAM 3’UTR (DC2); a bidirectional eEF1α promoter (DC3); a Cre60 coding sequence followed by the 3’ UTR of PbHSP70 (DC4); and a sequence corresponding to the 3’ UTR of PbDHFR (DC5). All elements of the DiCre plasmid were amplified by PCR using standard PCR conditions (using the CloneAmp HiFi PCR premix) and sequentially ligated (In-Fusion HD Cloning Kit, Clontech) into an acceptor plasmid containing a TgDHFR/TS cassette flanked by two LoxP sites. The DC2 and DC4 fragments were amplified using genomic DNA from DiCre-expressing *P. falciparum* parasites (kind gift of Kai Wengelnik and Mike Blackman). The resulting plasmid sequence was verified by Sanger sequencing (Eurofins Genomics) and linearized with *Sca*I and *Dra*III before transfection. The DC1 and DC5 fragments served as homology regions for double homologous recombination between the HSP70 promoter the PbDHFR 3’ UTR sequences flanking GFP at the modified p230p locus of PbGFP parasites. All primers used to construct the DiCre plasmid are listed in Table S1.

#### PbDCIII plasmid

The PbDCIII plasmid was designed to replace the mCherry cassette of the DiCre parental line with a cassette containing two fluorescent markers, GFP and ECFP, separated by a silent intron containing a LoxN site [11]. To permit the selection of transfected parasites, we also included a human dihydrofolate reductase marker (hDHFR) downstream of the GFP cassette, followed by the 3’ UTR of *P. berghei* calmodulin (CAM) gene. A 2A skip peptide was introduced between GFP and hDHFR to enable transcription of both cassettes under the control of a single HSP70 promoter. Furthermore, the GFP-2A-hDHFR-PbCAM cassette was floxed by two silent introns containing LoxN sites, as shown in **Figure 3A**. The LoxN-GFP-2A-hDHFR and LoxN-ECFP fragments were ordered as synthetic genes (Eurofins Genomics). All elements of the PbDCIII plasmid were amplified by PCR using standard PCR conditions (using the CloneAmp HiFi PCR premix) and sequentially ligated into the *SphI-SalI* restriction sites of a pUC18 vector (In-Fusion HD Cloning Kit, Clontech). The resulting plasmid sequence was verified by Sanger sequencing (Eurofins Genomics) and linearized with *NarI* before transfection. All primers used to construct the GFP-ECFP plasmid are listed in Table S1.

### Experimental animals, parasites and cell lines

Female SWISS mice (6–8 weeks old, from Janvier Labs) were used for all routine parasite infections. Parasite lines were maintained in mice through intraperitoneal injections of frozen parasite stocks and transmitted to *Anopheles stephensi* mosquitoes for experimental purposes. *P. berghei* sporozoites were isolated from infected female *Anopheles* mosquitoes and intravenously injected into C57BL/6J mice (female, 4-6 weeks old, Janvier Labs). A drop of blood from the tail was collected in 1ml PBS daily and used to monitor the parasitaemia by flow cytometry.

For parasite transfection, schizonts purified from an overnight culture of PbGFP or PbDiCre parasites were transfected with 5–10 μg of linearized plasmid by electroporation using the AMAXA Nucleofector device (program U033), as previously described [12], and immediately injected intravenously into the tail vein of SWISS mice. To permit the selection of resistant transgenic parasites, pyrimethamine (35 mg/L) was added to the drinking water and administered to mice, one day after transfection. The parasitaemia was monitored daily by flow cytometry and the mice sacrificed at a parasitaemia of 2-3%. The mice were bled and the infected blood collected for preparation of frozen stocks and isolation of parasites for genomic DNA extraction.

Pure transgenic parasite populations were isolated by flow cytometry-assisted sorting of mCherry or GFP-expressing blood stage parasites on a FACSAria II (Becton-Dickinson), as described [10]. Mice were injected intraperitoneally with frozen parasite stocks and monitored until the parasitaemia was between 0.1 and 1%. On the day of sorting, one drop of blood from the tail was collected in 1 ml PBS and used for sorting of 100 pRBCs which were recovered in 200 μl RPMI +20% foetal calf serum (FCS), and then injected intravenously into two mice (100 μl each). The mice were sacrificed at a parasitaemia of 2-3% and the blood recovered for preparation of frozen stocks and genotyping.

*Anopheles stephensi* mosquitoes were reared at 24-26°C with 80 % humidity and permitted to feed on infected mice that were anaesthetised, using standard methods of mosquito infection as previously described [13]. Post-feeding, *P. berghei*-infected mosquitoes were kept at 21°C and fed on a 10% sucrose solution. Salivary gland sporozoites were collected between 21 and 28 days post-feeding from infected mosquitoes, by hand dissection and homogenisation of isolated salivary glands in complete DMEM (DMEM supplemented with 10% FCS, 1% Penicillin-Streptomycin and 1% L-Glutamine).

HepG2 cells (ATCC HB-8065) were cultured in DMEM supplemented with 10% FCS, 1% Penicillin-Streptomycin and 1% L-Glutamine as previously described [14].

### Rapamycin treatment

DiCre recombinase mediated excision of targeted DNA sequences *in vivo* was achieved by oral administration of 200 μg Rapamycin (1mg/ml stock, Rapamune, Pfizer) to mice, 24 hours prior to transmission to mosquitoes. In order to achieve excision in the mosquito stages, 10 μg rapamycin (1 mg/ml stock solution in DMSO, Sigma-Aldrich) was added to 10 ml 10% sucrose solution and used to feed mosquitoes. The rapamycin dose was refreshed every alternate day along with the sucrose solution.

### Genotyping PCR

Infected mice were sacrificed at a parasitaemia of 2-3% and the infected blood collected and passed through a CF11 column (Whatman) to deplete leucocytes. The RBCs collected were then centrifuged and lysed with 0.2% saponin (Sigma) to recover parasite material for genomic DNA isolation using a kit (Qiagen DNA Easy Blood and Tissue Kit), according to the manufacturer’s instructions. Specific PCR primers were designed to check for wild-type and recombined loci and are listed in Table S2. All PCR reactions were carried out using Recombinant Taq DNA Polymerase (5U/μl from Thermo Scientific) and standard PCR cycling conditions.

### *In vitro* infections, immunofluorescence assays and microscopy

#### Infection of hepatocytes

HepG2 cells were seeded in collagen-coated culture plates, at a density of 30,000 cells/well in a 96-well plate for flow cytometry analysis or 100,000 cells/well in 8 well μ-slide (IBIDI) for immunofluorescence assays, 24 hours prior to infection with sporozoites. On the day of infection, the culture medium in the wells was refreshed with complete DMEM, followed by the addition of 10,000 sporozoites and incubation for 3 hours at 37°C. After 3 hours, the wells were washed twice with complete DMEM and then incubated for another 24-48 hours at 37°C and 5% CO2.

For quantification of EEF numbers, the cells were trypsinized after two washes with PBS, followed by addition of complete DMEM and one round of centrifugation. After discarding the supernatant, the cells were either directly re-suspended in FACS buffer (PBS + 3% FCS) for flow cytometry, or fixed with 2% PFA for 10 minutes, subsequently washed once with PBS and then re-suspended in PBS. Cells were then analyzed on a Guava EasyCyte 6/2L bench cytometer equipped with 488 nm and 532 nm lasers (Millipore).

#### Immunofluorescence assays

For immunofluorescence assays, the cells were washed twice with PBS, then fixed with 4% PFA for 10 minutes followed by two washes with PBS, quenching with glycine 0,1 M for 5 minutes, permeabilization with 1% Triton X-100 for 5 minutes before washes with PBS and blocking in PBS with 3% bovine serum albumin (BSA). Cells were then incubated for 1h with goat anti-UIS4 primary antibody (1:500, Sicgen), followed by donkey anti-goat Alexa Fluor 594 or Alexa Fluor 488 secondary antibody (1:1000, Life Technologies).

#### Fluorescence microscopy

Live samples such as blood-stages, midguts and salivary glands from infected mosquitoes, were mounted in PBS and visualised live using a fluorescence microscope (Zeiss Axio Observer.Z1 fluorescence microscope equipped with a LD Plan-Neofluar 403/0.6 Corr Ph2 M27 objective). Due to spectral overlap between GFP and CFP channels, we set the exposure time according to the positive control and maintained the same exposure for both excised and non-excised parasites, in order to allow comparisons. All images were processed with ImageJ for adjustment of contrast.

### RESULTS

#### Generation of a DiCre-expressing *P. berghei* line

To evaluate the DiCre system in *P. berghei*, we first generated a parasite line that stably and constitutively expresses both components of the Cre enzyme. We assembled a construct encoding the N-terminal Cre 59 (residues Thr19-Asn59) and C-terminal Cre 60 (Asn60-Asp343) portions of the Cre fused at their N-terminus to FKBP12 and FRB, respectively (**Figure 1A**). The two components were placed under control of the constitutive bidirectional promoter eEF1alpha, and followed by the 3’ untranslated region (UTR) from PfCAM and PbHSP70, respectively. In addition, the construct contained a mCherry cassette under the control of an inactive truncated fragment of HSP70 promoter, and a TgDHFR/TS pyrimethamine resistance cassette. The TgDHFR/TS cassette was flanked by two LoxP sites, to allow Cre-mediated excision and production of drug selectable marker-free parasites (**Figure 1A**). A sequence corresponding to the 3’ UTR of PbDHFR was included at the end of the construct, for homologous recombination at the modified *p230p* locus of GFP-expressing *P. berghei* ANKA parasites (PbGFP) [10] (**Figure 1A**).

**Figure 1.**
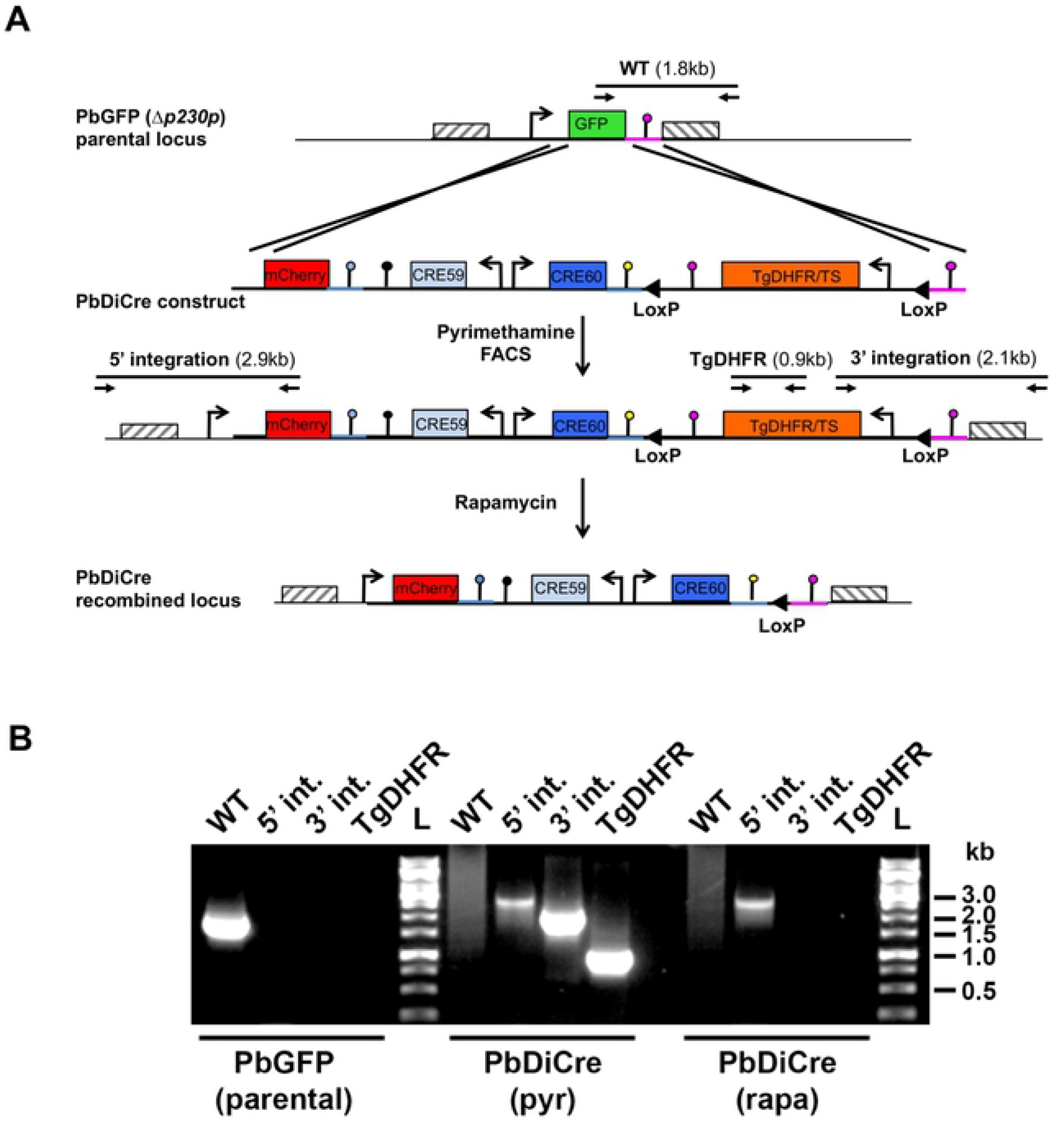
Generation of DiCre-expressing *P. berghei* parasites. **A.** Replacement strategy to modify the PbGFP locus for DiCre expression. The DiCre plasmid, containing the two Cre components in addition to a mCherry cassette and a TgDHFR/TS pyrimethamine resistance cassette, was integrated into a pre-modified P230p locus in *P.berghei*, where the P230p gene was previously replaced by a GFP cassette [10]. Following parasite transfection and selection with pyrimethamine, mCherry+ parasites were sorted by flow cytometry to exclude any residual GFP+ population. Genotyping primers and expected PCR fragments are indicated by arrows and lines, respectively. **B.** PCR analysis of genomic DNA isolated from parental PbGFP, pyrimethamine-selected (pyr) and rapamycin-treated (rapa) PbDiCre parasites. Confirmation of the predicted recombination events was achieved with primer combinations specific for WT, 5’ or 3’ integration. The TgDHFR primer combination was used to confirm the loss of the TgDHFR/TS cassette following rapamycin treatment.

Following transfection of PbGFP parasites, integration of the construct by double homologous recombination resulted in the replacement of GFP by mCherry and reconstitution of a functional HSP70 promoter to drive mCherry expression, along with insertion of the DiCre and TgDHFR/TS cassettes. Transfected parasites were selected with pyrimethamine and mCherry-positive parasites were sorted by FACS. The resulting parasite population was exposed to a single dose of rapamycin that was administered orally to mice. This treatment resulted in excision of the TgDHFR/TS cassette, as demonstrated by PCR genotyping (**Figure 1B**). Cloning by limiting dilution resulted in the final selectable-marker free mCherry-expressing PbDiCre parasite line.

#### PbDiCre parasites progress normally across the parasite life cycle

To exclude any interference of the DiCre cassette with parasite life cycle progression, we examined if these parasites could be transmitted to mosquitoes and back to mice. PbDiCre parasites formed oocysts in the midgut of infected mosquitoes and developed into sporozoites that colonized the insect salivary glands (**Figure 2A**). Similar numbers of salivary gland sporozoites were recovered from mosquitoes infected with PbDiCre and parental PbGFP parasites (**Figure 2B**), confirming that PbDiCre parasites develop normally in the mosquito. In addition, when incubated with a monolayer of HepG2 cells, PbDiCre sporozoites formed similar numbers of exo-erythrocytic forms (EEFs) as PbGFP parasites (**Figure 2C**). Liver stage development of PbDiCre parasites was comparable to PbGFP, as evidenced by EEF size (**Figure 2D**) and formation of a UIS4-labeled PVM (**Figure 2E**). Finally, we injected PbDiCre and PbGFP sporozoites into C57BL/6J mice and observed no difference in either pre-patency or blood-stage parasitaemia between PbGFP-and PbDiCre-infected mice (**Figure 2F**), showing that PbDiCre parasites are capable of completing the parasite life cycle, similarly to the parental PbGFP parasites. In summary, these results confirmed that parasites constitutively expressing the DiCre cassette progress normally through the parasite life cycle, and can thus be used as a parental strain to target genes of interest.

**Figure 2.**
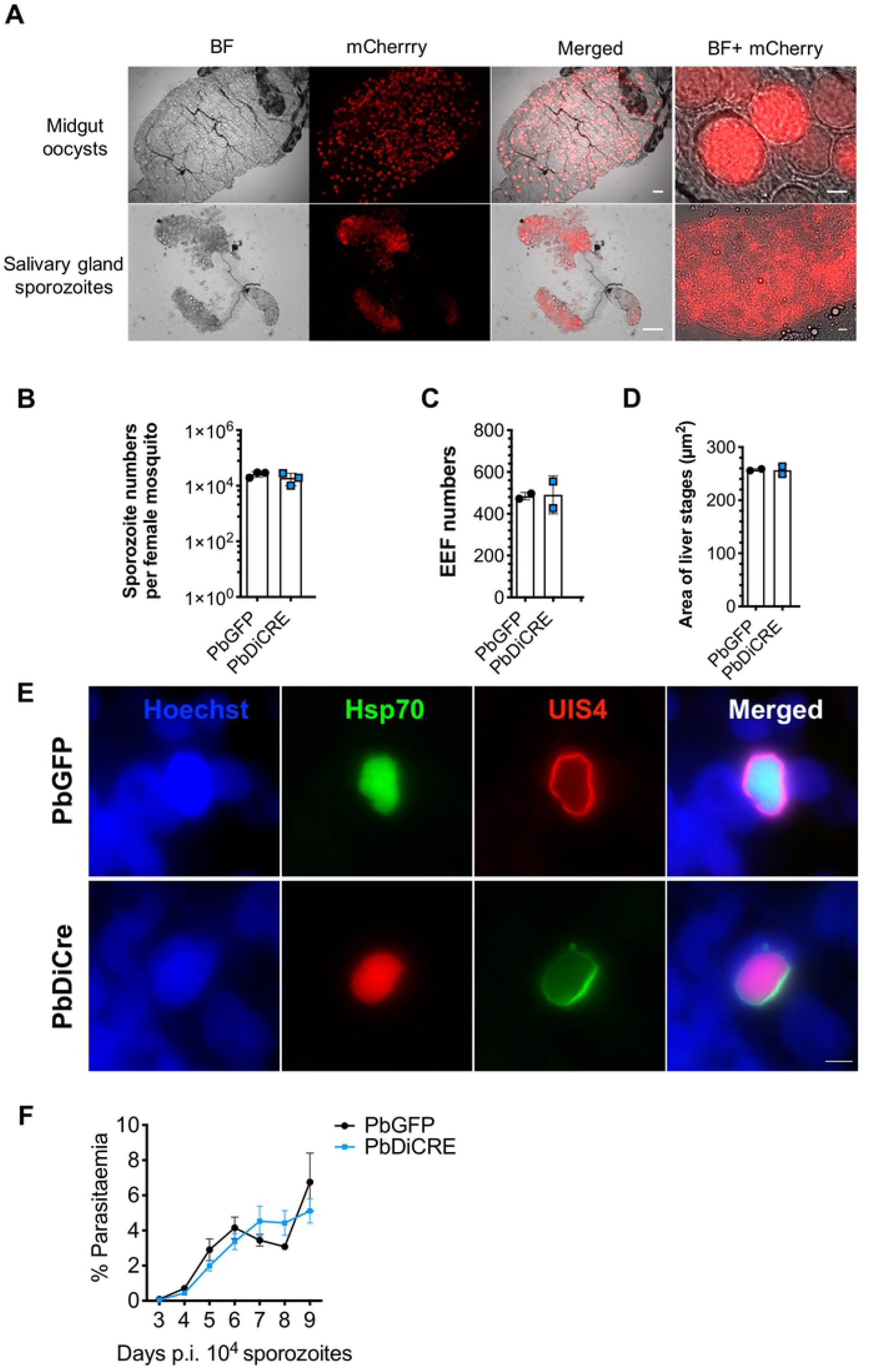
PbDiCre parasites show no defect in mosquito and liver stages. **A.** Fluorescence microscopy imaging of unfixed midgut oocysts and salivary glands from PbDiCre-infected mosquitoes. Scale bar, 100 μm for the midguts and salivary glands, and 10 μm for the magnified images. **B.** Comparison of sporozoite numbers isolated from salivary glands of female mosquitoes infected with PbGFP or PbDiCre parasites. Data shown are mean +/− SD of three independent experiments. **C.** Quantification of EEFs in HepG2 cell cultures infected with PbGFP or PbDiCre sporozoites. Data shown are mean +/− SD of two independent experiments. **D.** Quantification of EEF size (area) in HepG2 cell cultures infected with PbGFP or PbDiCre sporozoites. Data shown are mean +/− SD of two independent experiments. **E.** Images of PbGFP and PbDiCre EEFs in HepG2 infected cell cultures, 48 hours post-infection. Scale bar, 10 μm. **F.** Comparison of parasite development *in vivo* post-sporozoite injection. C57BL6/J mice (n = 5) were injected intravenously with 1 x 10^4^ PbGFP or PbDiCre sporozoites. Parasitemia was then followed daily by FACS. The data shown are mean +/− SEM of 5 mice per group.

#### Exposure to rapamycin during the mammalian blood stages is efficient to generate recombined mosquito stages

In order to assess the efficacy and robustness of the DiCre system to target genes of interest during parasite transmission to the mosquito, we designed a reporter construct that allows a simple assessment of site-specific rapamycin-induced DNA excision, using fluorescent markers. The designed construct replaced the endogenous mCherry cassette in PbDiCre parasites with a gene sequence comprised of GFP-2A-hDHFR-PbCAMutr flanked by LoxN sites, and placed under the inactive truncated fragment of HSP70 promoter (**Figure 3A**). An additional eCFP cassette lacking a promoter was inserted downstream of the second LoxN site (**Figure 3A**) thereby permitting selective expression of GFP or CFP before or after rapamycin-induced excision, respectively. Proper integration of the construct was confirmed by genotyping PCR (**Figure 3B**), resulting in a PbDCIII parasite line that provided us with a suitable tool to visually monitor rapamycin-induced excision in all developmental stages of the parasite.

**Figure 3.**
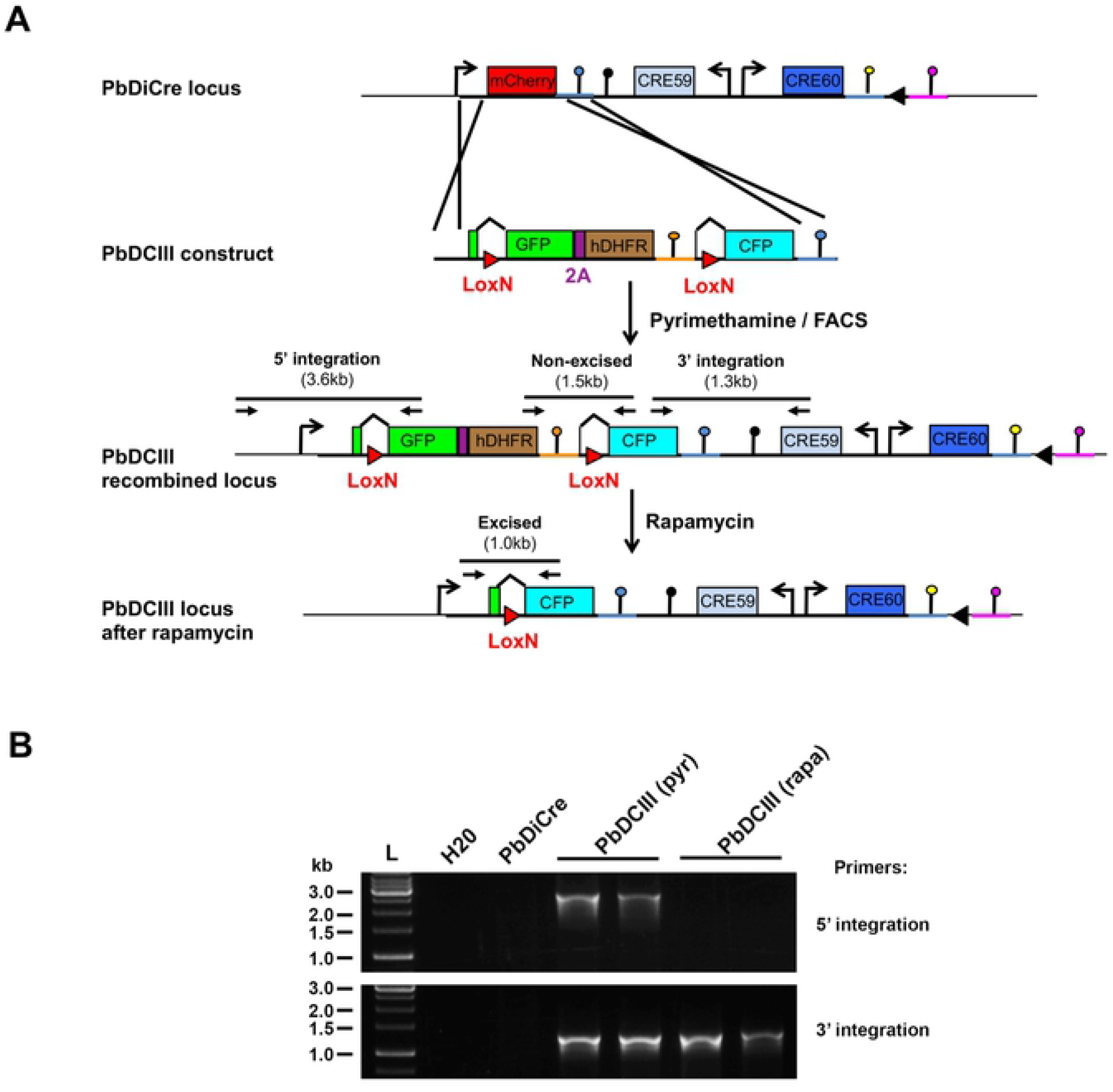
Generation of a PbDCIII reporter line to monitor Cre activity. **A**. Replacement strategy to modify the mCherry locus in PbDiCre parasites and generate the PbDCIII parasite line. The DCIII construct contains a GFP-hDHFR cassette flanked by LoxN sites, and followed by a CFP cassette. Integration of the construct results in GFP expression instead of mCherry. Following rapamycin treatment, Cre-mediated recombination results in excision of the GFP cassette and expression of CFP. Genotyping primers and expected PCR fragments are indicated by arrows and lines, respectively. **B.** PCR analysis of genomic DNA isolated from parental PbDiCre, pyrimethamine-selected (pyr) and rapamycin-treated (rapa) PbDCIII parasites. Confirmation of the predicted recombination events was achieved with primer combinations specific for 5’ and 3’ integration.

In order to verify that exposure of blood-stages to rapamycin results in efficient gene excision, we infected mice with PbDCIII parasitised red blood cells (pRBCs) and treated them with one dose of rapamycin. Untreated mice were used as controls. At 48h post-treatment, direct examination by fluorescence microscopy confirmed the presence of CFP-positive (CFP+) parasites in the blood of rapamycin-treated mice (**Figure 4A**), while only GFP-positive (GFP+) parasites were observed in untreated mice, suggesting that gene excision in blood-stages was very efficient after rapamycin treatment. Although genotyping PCR confirmed complete excision in rapamycin-treated PbDCIII parasites, excision bands could also be amplified by PCR in untreated PbDCIII parasites (**Figure 4B**), suggesting some level of activity of the Cre in the absence of rapamycin.

**Figure 4.**
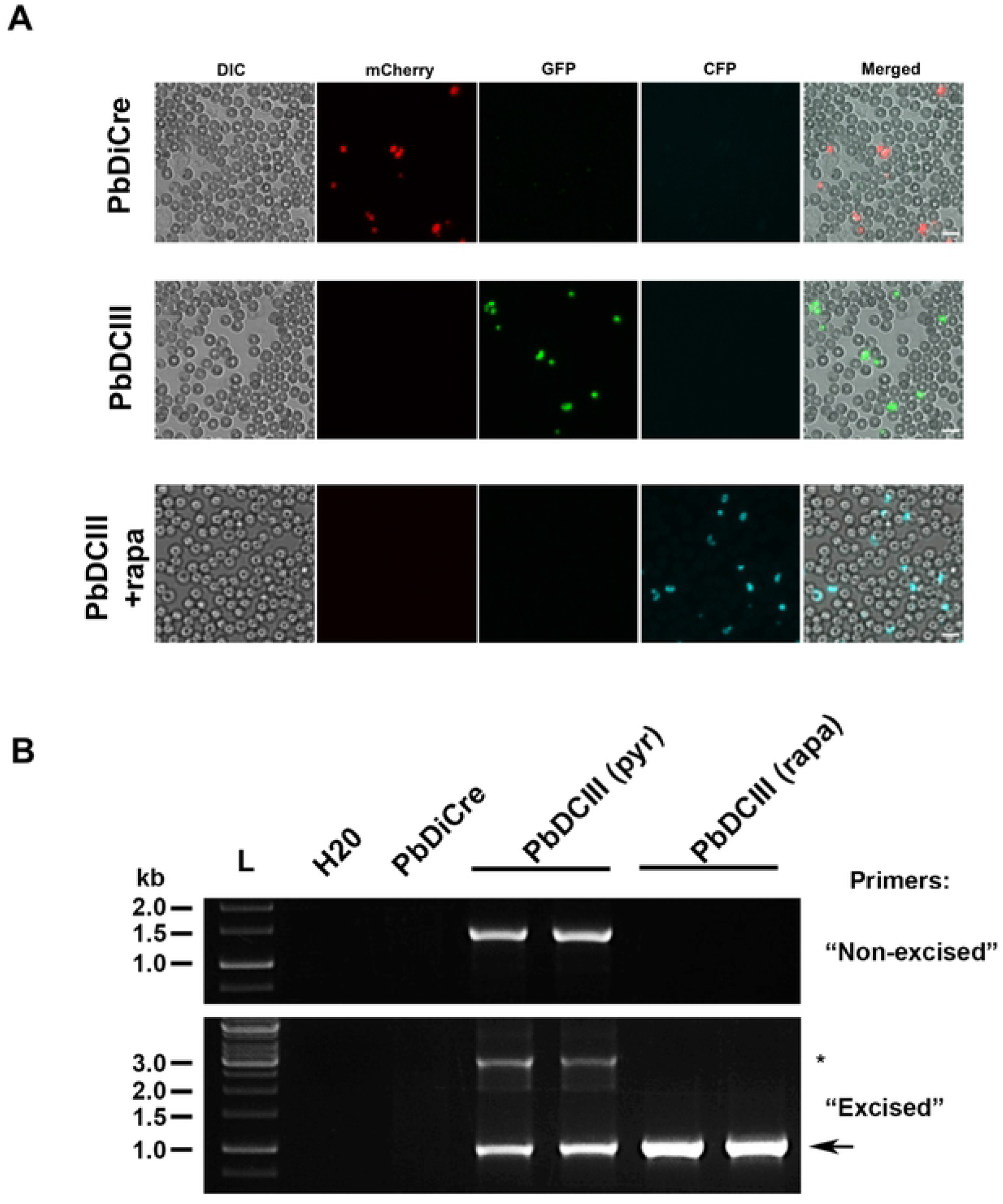
Exposure to rapamycin during the mammalian blood stages. **A.** Fluorescence microscopy of unfixed blood stages in the parental PbDiCre parasites (upper panels) and in PbDCIII parasites before (middle panels) and after (lower panels) rapamycin treatment. Scale bar, 10 μm. **B.** PCR analysis of genomic DNA isolated from parental PbDiCre, pyrimethamine-selected (pyr) and rapamycin-treated (rapa) PbDCIII parasites, using primer combinations specific for non-excised or excised locus. A band corresponding to a non-excised locus can be amplified with the “excised” primer combination and is indicated with an asterisk.

In the next step, we evaluated if DNA excision could also be induced in sexually committed parasites/gametocytes. Mice infected with PbDCIII parasites were administered rapamycin 24 hours prior to transmission to mosquitoes (**Figure 5A**), following which infection and development of both treated and untreated parasites in the mosquito was monitored by fluorescence microscopy. We predominantly observed CFP+ oocysts and salivary gland sporozoites in mosquitoes infected with rapamycin-treated PbDCIII parasites, showing that one dose of rapamycin is sufficient to induce efficient excision in transmission stages (**Figure 5B**).

**Figure 5.**
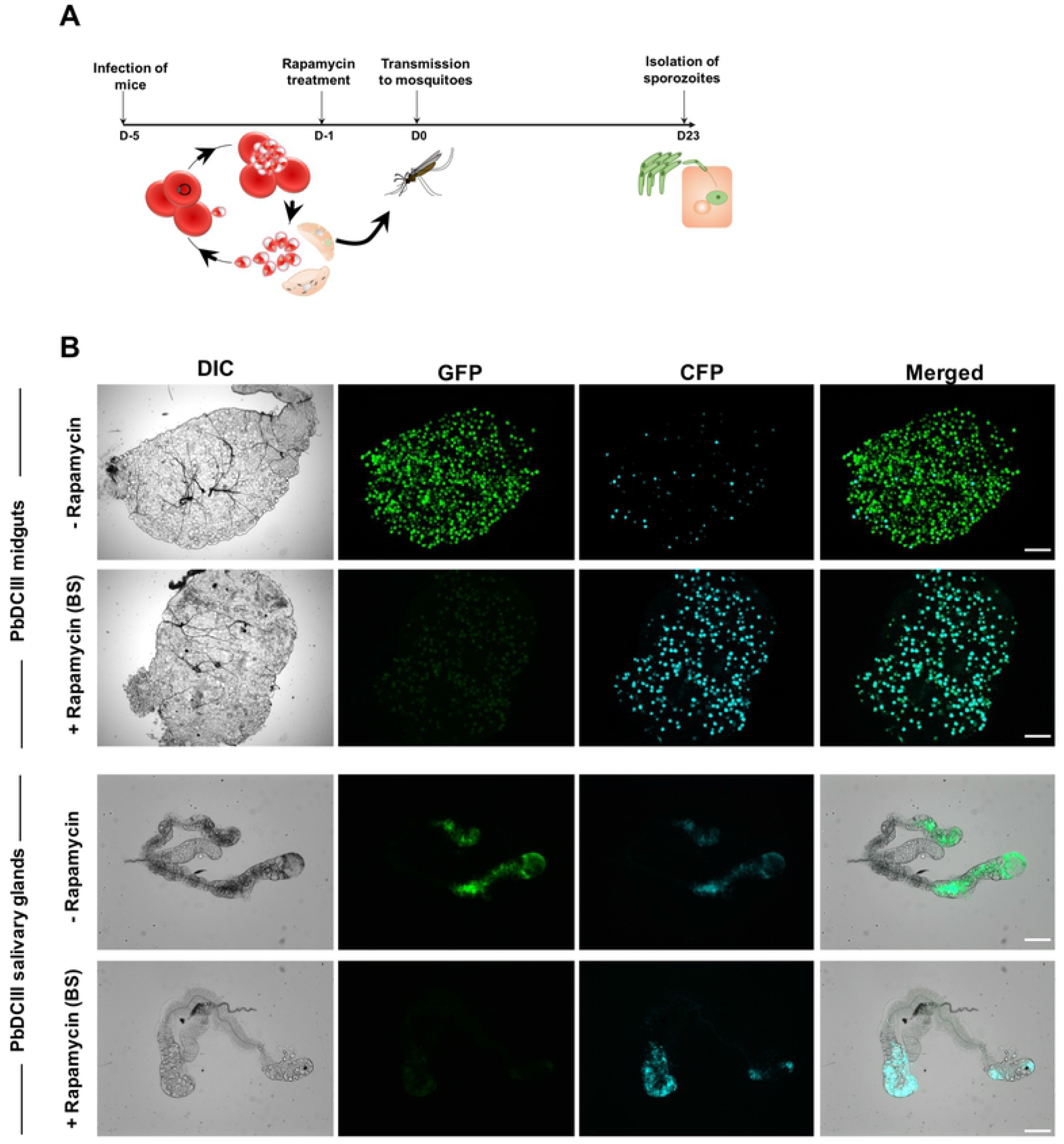
Rapamycin-mediated DNA recombination in sexual blood stages. **A.** Illustration of the rapamycin treatment protocol applied to sexually committed blood stages prior transmission to mosquitoes. **B.** Fluorescence microscopy of unfixed midguts and salivary glands isolated from PbDCIII-infected mosquitoes parasites, before (-rapamycin) or after blood-stage rapamycin treatment (+ rapamycin BS). Scale bar, 200 μm for midguts and 100 μm for the salivary glands.

#### Exposure of mosquito stages to rapamycin allows gene deletion in sporozoites

Due to optimal activity of the Cre enzyme at 37 °C, the DiCre system has consistently been used to investigate the function of genes in *Plasmodium* blood-stages, but lacks applicability when it comes to studying the function of genes involved in ookinete formation or establishment of infection in the insect vector. In order to overcome this drawback, we tested if exposure to rapamycin of infected mosquitoes, instead of blood-stages, would result in efficient DNA excision. For this purpose, we fed PbDCIII-infected mosquitoes on a sucrose solution containing 1 μg/ml rapamycin, for two weeks, starting at day 5 post-transmission (**Figure 6A**). Interestingly, both oocysts and salivary glands isolated from treated mosquitoes were CFP+, showing that DiCre-mediated DNA excision can also be induced by treating mosquitoes with rapamycin (**Figure 6B**). In these experiments, rapamycin exposure during parasite development in the mosquito was associated with a reduction of the number of salivary gland sporozoites (**Figure 6C**). However, a similar reduction was observed when rapamycin was administered to mice prior to mosquito blood feeding (**Figure 6C**). With both protocols, sporozoites remained infectious to HepG2 cells cultures (**Figure 6D**), and both rapamycin treatment regimens resulted in almost complete parasite DNA recombination as revealed by the switch from GFP to CFP expression (**Figure 6E**).

**Figure 6.**
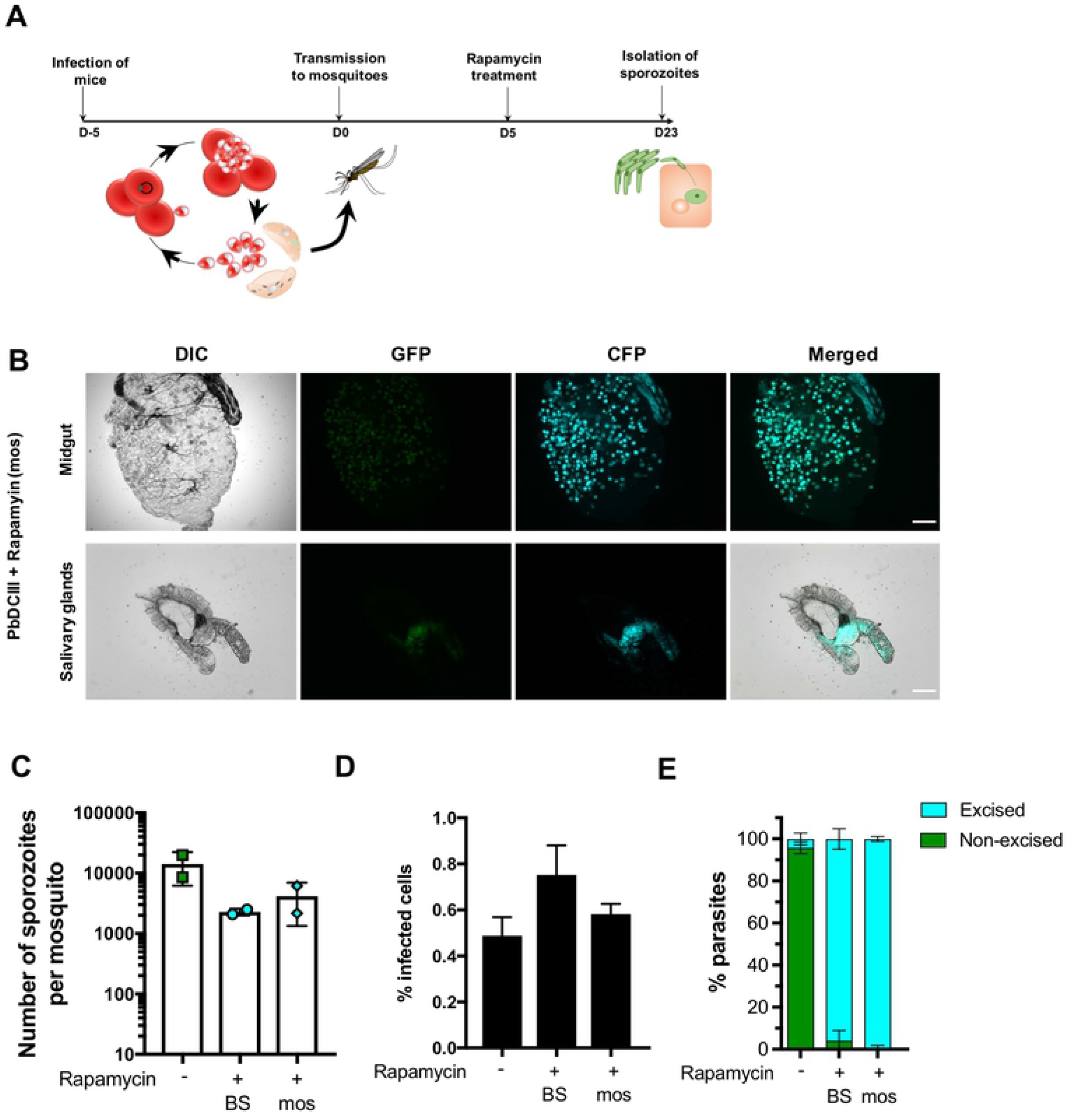
Rapamycin-mediated DNA recombination in mosquito stages. **A.** Illustration of the rapamycin treatment protocol applied to infected mosquitoes. **B.** Fluorescence microscopy of unfixed midguts and salivary glands isolated from PbDCIII-infected rapamycin-treated mosquitoes. Scale bar, 200 μm for midguts, and 100 μm for the salivary glands. **C.** Quantification of salivary gland sporozoites in mosquitoes infected with the PbDCIII parasites, after rapamycin exposure of mice (BS) or mosquitoes (mos) in comparison to the untreated parasites. Results shown are mean+/− SD of two independent experiments. **E.** Quantification of infected cells in HepG2 cell cultures infected with the PbDCIII parasite line, after rapamycin exposure of mice (BS) or mosquitoes (mos) in comparison to untreated control. Results shown are mean+/− SD of five technical replicates and are representative of two independent experiments. **F**. Quantification of excised CFP+ and non-excised GFP+ EEFs *in vitro* in infected HepG2 cell cultures, as determined by FACS. Results shown are mean+/− SD of five technical replicates and are representative of two independent experiments.

Overall, these results showed that the DiCre system can be used to target genes in different developmental stages both in the mammalian host and in the mosquito vector.

## DISCUSSION

In spite of numerous advances in gene editing technologies that are currently available to investigate the function of *Plasmodium* genes, the role of blood-stage essential genes in pre-erythrocytic developmental stages remains poorly explored. We implemented the DiCre system in *P. berghei* parasites and demonstrate how the system can be used to conditionally delete genes in *P. berghei* mosquito stages. We first generated a marker-free fluorescent parasite line in *P. berghei*, which constitutively expresses both components of the dimerisable Cre recombinase [9], by introducing components of the DiCre along with an mCherry cassette into a non-essential P230p locus [15]. Asexual development, sporogony in the mosquito, sporozoite infectivity in mice and EEF development *in vitro* were comparable to a reference PbGFP line [10], showing that the components of the DiCre did not interfere with parasite development or transmission. We then confirmed the integrity of the Cre components in inducing excision after rapamycin treatment, using the PbDCIII reporter line. The data obtained with the PbDCIII parasite line not only validated the activity of the Cre enzyme in PbDiCre, but also confirmed that a single dose of rapamycin was sufficient to induce close to 100% excision in blood-stages, consistent with previously published data [2,16]. Excision events were also noted in untreated PbDCIII parasites by PCR and in a minor proportion of oocysts and sporozoites, suggesting some leakiness in a minor proportion of parasites. Alternatively, we cannot exclude that co-housing of untreated and rapamycin-treated mice may result in the exposure of untreated mice to rapamycin excreted in the faeces of treated mice. Nevertheless, rapamycin-induced DNA recombination in gametocytes was very efficient as observed by the lack of GFP+ oocysts in the mosquito, suggesting that targeted deletion of blood-stage essential genes can successfully be achieved in sexual stages.

A recent study used a *P. falciparum* NF54 DiCre parasite line to generate conditional knockouts of AMA1 and FIKK7.1 genes, and reported a significant reduction in oocyst numbers after transmission of rapamycin treated parasites to mosquitoes [17]. The authors speculated that treatment with rapamycin was mainly responsible for the reduction they observed. In agreement with these observations, we observed a reduction of salivary gland sporozoite numbers after rapamycin treatment in the DCIII parasite line. In light of these findings, we sought to circumvent the effects of rapamycin on oocyst formation by treating mosquitoes with rapamycin instead of blood-stages. Mosquitoes infected with untreated PbDCIII parasites were fed with rapamycin 5 days after the blood-meal, to avoid any effects on ookinetes and the formation of oocysts. However, a similar reduction of the number of salivary gland sporozoites was observed irrespective of the rapamycin treatment regimen. This detrimental effect could be due to inhibition of the target of rapamycin (TOR) pathway, which is known to play a role in nutritional sensing in the mosquito [18], and may participate in oocyst formation and/or development.

When rapamycin was administered to infected mosquitoes, we observed highly efficient excision of the GFP in PbDCIII parasites, and we only observed excised CFP+ EEFs in infected hepatocytes. Thus, rapamycin treatment can successfully be adapted to different stages in the mosquito, such as treatment of midgut sporozoites prior to salivary gland invasion or treatment of salivary gland sporozoites before hepatocyte invasion, although we predict that the percentage of excision might vary with shorter duration times.

In summary, we generated a fluorescent PbDiCre line and show that it can be used to investigate the function of essential blood-stage genes in mosquito stages. By treating parasites with rapamycin in the blood or in the mosquito, we demonstrate the versatility of the DiCre system in targeting genes of interest at multiple stages of development. Overall, our study introduces a new robust and versatile methodology to conditionally delete genes in different stages of *Plasmodium* parasites, which should help deciphering the function of essential genes across the parasite life cycle.

## ACKNOWLEDGMENTS

The authors would like to thank Jean-François Franetich, Maurel Tefit, Mariem Choura and Thierry Houpert for rearing of mosquitoes. DNA encoding the Cre elements was a kind gift from Kai Wengelnik and Mike Blackman. The research leading to these results has received funding from the Laboratoire d’Excellence ParaFrap (ANR-11-LABX-0024), and the Agence Nationale de la Recherche (ANR-16-CE15-0004 and ANR-16-CE15-0010). ML was supported by a ‘DIM1Health’ doctoral fellowship awarded by the Conseil Régional d’Ile-de-France.

